# Isolation of a novel plant growth-promoting *Dyella* sp. from a Danish natural soil

**DOI:** 10.1101/2025.01.24.634660

**Authors:** Laura Dethier, J. Rasmus P. Jespersen, Jemma Lloyd, Elena Pupi, Wanru Zhou, Fang Liu, Yang Bai, Barbara Ann Halkier, Deyang Xu

**Author notes:** NIRAS A/S, Sortemosevej 19, 3450 Allerød, Denmark (J. Rasmus P. Jespersen). Fondazione Edmund Mach, Via E. Mach 1, 38098 San Michele All’Adige TN, Italy. Funding DISTINGUISHED INVESTIGATOR from Novo Nordisk foundation (NNF20OC0060824 to B.A.H), National Key Research and Development Projects (2022YFF1001800 to Y. B.) and Science Xplore Award (XPLORER-2023-1017 to Y.B.).

## Abstract

Environmental soils are natural reservoirs of unexplored microbes, including potentially beneficial microbes to improve plant performance. Here, we isolated 75 bacterial strains from surface-sterilized roots of *Arabidopsis thaliana* (Arabidopsis) grown in a natural soil derived from an alder swamp. Culture-dependent isolation of individual strains from the roots followed by monoassociation-based screening identified seven bacteria that promoted Arabidopsis seedling weight. Of those, we identified a new species from the *Dyella* genus which increased biomass of tomato and Arabidopsis seedlings in agar as well as the shoot biomass of Arabidopsis grown in both the alder swamp soil and potting soil. *Dyella sp*. A4 specifically promoted the elongation of lateral roots without affecting lateral root number and primary root elongation. The new *Dyella sp*. A4 expands the toolbox of biostimulants for plant growth promotion via modulating root architecture.

## Introduction

Soils are inhabited by diverse microbial communities that vary in composition across different geographic locations, dependent on factors such as pH, nutrient availability, texture and moisture content (Fierer and Jackson, 2006; Karimi et al., 2018; Xia et al., 2020). From the pool of microbes present in bulk soils, a subset is recruited to establish close associations with plant roots – collectively known as the root microbiota – resulting in a range of beneficial, neutral or pathogenic interactions with the host plant (Lundberg et al., 2012). The potential to harness beneficial members of the root microbiota to improve plant growth, nutrient acquisition and disease resistance has received considerable interest and investment in the last decade (Finkel et al., 2017; Zhang et al., 2019).

Natural soils represent a repository of beneficial microbes to be explored, and research has focused on the isolation of microbes from nutrient-poor and/or arid soils that can promote plant resilience to abiotic stresses (Eida et al., 2018; Khan et al., 2023). For example, the bacterium *Isoptericola* sp. AK164 isolated from the rhizosphere of *Avicennia marina* plants grown in the red sea coast, improved Arabidopsis biomass (Alghamdi et al., 2023). Another study found that the root fungal endophyte F229, isolated from *Arabis alpina* grown in a natural P-limited soil, promoted plant growth and shoot P accumulation, pointing towards microbial partners in Brassicaceae species that are incapable of forming symbiosis with AM fungi (Almario et al., 2017). Root architecture is important for plant nutrient uptake and some microbes can modify aspects of root architecture to promote plant growth (Khoso et al., 2024). For instance, the rhizobacterium *Bacillus megaterium* strain WW1211 promoted lateral root initiation and shoot biomass by inducing auxin biosynthesis, redistribution and signaling in Arabidopsis (Wang et al., 2021). Additionally, bacterial strains that modulate Arabidopsis root branching through ethylene or cytokinin signaling rather than auxin signaling, have been identified (Gonin et al., 2023; López-Bucio et al., 2007; Ortíz-Castro et al., 2008). In this study, we isolated different bacteria from the root endosphere of Arabidopsis grown in a Danish natural alder swamp soil and identified a novel plant growth-promoting (PGP) *Dyella* sp. A4 that specifically promotes lateral root elongation, without affecting primary root length and lateral root number.

## Materials and Methods

### Soil collection and processing

The soil used in this study was collected from an alder swamp (55°58’05.6”N, 12°16’16.7”E) within Strødam Nature Reserve in Denmark in March 2019 with permission obtained from Strødam committee. The site is annotated in Biowide (Biodiversity in Width and Depth) project with a site ID 82 (Brunbjerg et al., 2019). The soil is a wet, organic soil and the vegetation at the site consists of alder (*Alnus glutinosa*), birch (*Betula sp*.) and sweet violet (*Viola odorata*). The top 30 cm of soil was collected to a total of 100 kg, air dried and sieved (10-mm sieve) to remove leaf litter and rocks. Approximately half of the soil was distributed into 3 kg plastic bags (300×600×0.08) and hermetically sealed before sterilization by gamma irradiation (18 kGy on each side of the soil bag; Sterigenics, Espergaerde, Denmark) to eliminate microbes. The gamma irradiation-sterilized bags were kept sealed until use and any leftover soil from an opened bag was discarded. Sterility was verified by suspending 5 g of sterilized and non-sterilized soil in 15 ml sterile water, shaking for 2 h and plating the supernatant onto nutrient agar media [0.5% peptone, 0.5% bacto yeast extract, 0.5% NaCl, 1.5% g Bacto agar (BD Difco)] in triplicates. No microbial growth was observed on the nutrient agar plates streaked with the sterilized soil sample (Figure S1A). All bags containing soil were stored at 4°C until further experiments.

### Plant materials and growth conditions

Arabidopsis Col-0 seeds were laboratory stocks, while DR5rev::GFP (CS9361) seeds were obtained from the Arabidopsis Biological Resource Center (ABRC). Plants were grown under long-day conditions (16 h light, 140 μmol m^−²^ s^−¹^, 22°C, 55–60% relative humidity) in a growth chamber. Tomato seeds (*Solanum lycopersicum* cv. Moneymaker) obtained from Frøsnapperen were grown under 16 h light, 100 μmol m^−²^ s^−¹^, 25°C and 55% relative humidity.

### Culture-independent analysis of the alder swamp soil and root-associated microbiota

We profiled bacterial communities from unplanted bulk soil and plant roots (endosphere and rhizoplane) of plants grown in alder swamp soil. Arabidopsis seeds were sown in alder swamp soil mixed with sand (1:1 v/v) and grown for 5 weeks. Roots were washed in phosphate buffered saline (PBS), placed in Lysing Matrix E tubes (MP Biomedicals), and stored at -80°C with unplanted soil aliquots. The experiment was conducted twice with 7 root and 3 soil replicates. DNA was isolated with the FastDNA Spin Kit for Soil (MP Biomedicals), diluted to 3.5 ng/μl, and amplified in a two-step PCR targeting the 16S rRNA V5–V7 region (799F-1193R) (Zhang et al., 2019). The first PCR (25 cycles) and second PCR (8 cycles) were performed in triplicate reactions in a 30 μl reaction volume containing 3 µl of diluted DNA, 0.75 U PrimeSTAR HS DNA polymerase, 1x PrimSTAR buffer (Takara), 0.2 mM dNTPs and 10 pM barcoded primers. PCR products were pooled and verified by agarose gel electrophoresis. Correct-size bands were excised, purified with the PCR Clean-Up System (Promega) and quantified. Amplicon libraries were pooled (200 ng each), purified, and sequenced on the Illumina HiSeq 2500 platform using 1.2 μg of library.

### Isolation of bacteria from the root endosphere

Arabidopsis seeds were surface-sterilized in 70% ethanol for 20 min, rinsed with sterile water, and stratified at 4°C in the dark. Seeds were sown in pots containing alder swamp soil mixed with sand (1:3 v/v) and grown in a growth chamber. Bacteria were isolated from eight-week-old plants using a modified limiting dilution method (Zhang et al., 2021). Roots were washed in deionized water to remove soil and incubated in PBS (180 rpm, 15 min, 24°C) before plating the last wash onto 0.5x tryptic soy broth (TSB; Sigma-Aldrich) agar. For endophytes, roots were sterilized with 0.25% sodium hypochlorite (30 s), rinsed 3 times with sterile water, and dried on sterile filter paper. Sterilization was verified by plating the final rinse, with no microbial growth observed after 7 days at 24°C (Figure S1B). Sterilized root tissue (20 mg) was homogenized in 200 µl sterile 10 mM MgCl_2_, incubated in MgCl_2_ for 15 min, and transferred to 10x TSB for serial dilutions (222x, 148x, and 74x). Each dilution was distributed into fifteen 96-well plates (Sarstedt), using 3 control plates containing sterile TSB. Plates were incubated in the dark at 24°C. After 2 weeks, approximately 20% bacterial growth was observed in 74x dilution plates, and isolates were preserved as 40% glycerol stocks at -80°C. Purification onto 0.5x TSB agar yielded 290 isolates, with 3.6% of wells containing a mixture of isolates.

### Identification of bacterial isolates and phylogenetic analysis

The 16S rRNA gene of isolates was amplified using primers 27F and 1492R (Zhang et al., 2021). Bacterial colonies were suspended in 100 µl water, and 1 µl was added to a 24 µl PCR reaction mix containing 2.5 µl 10x buffer, 1 µl 50 mM MgCl_2_, 2 µl 2.5 mM dNTPs, 0.75 µl 10 µM primers, 0.6 µl Taq polymerase, and 16.4 µl water. PCR cycling consisted of 94°C for 2 min, 30 cycles of 94°C for 30 s, 55°C for 30 s, 72°C for 2 min, and a final elongation at 72°C for 10 min. PCR products were loaded on 1% agarose gel, excised, and purified using the E.Z.N.A. Gel Extraction Kit (Omega Bio-tek), then sequenced via Sanger sequencing. Low-quality ends were trimmed, and sequences were aligned and searched against GenBank using BLASTN (Tables S6 and S7). Final isolates were stored at -80°C as 40% glycerol stocks. Phylogenetic relationships were inferred by aligning sequences with MUSCLE 5.1 (Edgar, 2004) and constructing a maximum likelihood tree in PhyML 3.3.2 (Guindon et al., 2010) using a GTR model and 1,000 bootstrap replicates. The consensus tree was visualized and edited in iTOL (Letunic and Bork, 2021).

### Inoculation of Arabidopsis with all isolates separately in agar

Isolates were inoculated into agar following gnotobiotic growth protocols (Ma et al., 2022). Arabidopsis seeds were sown on 0.5x Murashige and Skoog (MS) agar (0.5x MS with vitamins (Duchefa), 0.5% sucrose, 1% Bacto agar, pH 5.7), stratified in the dark at 4°C for 2 days, and germinated for 6 days. Seedlings were then transferred to new 0.5x MS agar plates (1% Bacto agar, pH 5.7) inoculated with individual bacterial strains or mock-treated controls. For the inoculations, strains were streaked from glycerol stocks onto 0.5x TSB agar and incubated at 28°C. Bacterial cells were then grown in 0.5x TSB medium (28°C, 200 rpm) for 1-14 days, pelleted, washed in 10 mM MgCl_2_, and inoculated into warm 0.5x MS agar at a final OD600 of 0.0005. For controls, 10 mM MgCl_2_ was used. The agar was poured into square plates and eight 6-day-old seedlings were transferred to each plate, with 2-3 plates per treatment. Seedlings were grown for an additional 10 days before measuring fresh weight.

### Whole-genome sequencing of *Dyella* sp. A4 and annotation

DNA was extracted from an overnight culture of *Dyella* sp. A4 using the FastDNA SPIN Kit for Soil (MP Biomedicals), followed by ethanol precipitation. DNA yield and quality were assessed using the NanoDrop 2000. Library preparation and sequencing were conducted by Novogene (Cambridge, UK) with the Illumina NovaSeq 6000 platform, generating ∼9 million 150 bp paired-end reads. Low quality reads and adapters were filtered off using Trimmomatic (v.0.33) (Bolger et al., 2014) under SLIDINGWINDOW:4:20 MINLEN:50. Contigs were assembled from clean reads using Spades (version 3.15.5) in isolate mode, resulting in 124 contigs with an average length of 40,442 bp. Genome quality was assessed using CheckM (v.1.2.1) for completeness and contamination, and taxonomy was assigned using GTDB-tk (v.2.0.0) based on R207 GTDB genome database. To infer the novelty of this species, we calculated pairwise distance between *Dyella* sp. A4 and the bacterial genome in R207 GTDB database using Mash (2.3). When the shortest distance between *Dyella* sp. A4 versus GTDB genome is larger than 5%, this species is defined as novel species, which is the case for A4 with the shortest Mash distance equal to 0.076. To infer the gene content and functional potential encoded within the strain, we predicted the coding sequence using prodigal (v.2.6.3) under single genome mode by setting -p to single. Functional characterization of all predicted genes was performed by blast against the KEGG database (2021 April version) using diamond v2.0.15 under the sensitive mode, and the e-value cutoff was set to 1.0e-5. To ensure the accuracy of the annotated function, only hits with identity larger than 50% were kept. Based on the KEGG gene and KO mapping information provided by the KEGG database, the annotated genes were aggregated to KEGG orthologs (KO). To infer the plant growth-promoting potential encoded by the strain, we curated a list of PGP traits based on published articles, including functions related to nutrient release, hormone biosynthesis and stress tolerance along with their KEGG ortholog information (Table S5). These PGP traits were then summarized as present or absent within *Dyella* sp. A4 by linking gene annotation results and the curated PGR ortholog information.

### Inoculation of Arabidopsis and tomato with *Dyella* sp. A4 in agar

*Dyella* sp. A4 was inoculated into 0.5x MS agar at a final OD600 of 0.0005, and 6 to 8 six-day-old Arabidopsis seedlings were transferred as described above. Primary root elongation was tracked by marking root positions on the plate surface post-transfer. Plates were imaged at 4, 6 and 8 dpi, and root elongation, lateral root number, and lateral root length were measured using EZ-Rhizo software (Armengaud et al., 2009). For confocal microscopy of the DR5rev::GFP reporter line, roots grown on axenic and *Dyella* sp. A4-inoculated MS plates were imaged with a Leica SP5 X confocal microscope (60x objective). Roots were stained with propidium iodide and mounted in water for imaging. Propidium iodide was excited at 561 nm with emissions detected at 570-630 nm, while GFP was excited at 488 nm with emissions detected at 500-530 nm. Imaging was performed at 2 and 6 dpi. For *Dyella* sp. A4 inoculation of tomato seedlings, seeds were sterilized (70% ethanol for 1 min, 0.5% sodium hypochlorite for 20 min, followed by rinses in water) and sown on 0.5x MS agar (without sucrose). Seeds were incubated in the dark at 24°C for 3 days. *Dyella* sp. A4 was inoculated into 100 ml of 0.5x MS agar at a final OD600 of 0.000125, which was poured into plant tissue culture boxes (95 × 95 × 105 mm). As a mock control, 100 ml of 0.5x MS agar was supplemented with MgCl_2_. Three to four germinated seedlings were transferred to each box. Three independent experiments were performed, with 3-4 boxes per treatment. Shoot fresh weight was measured after 24 days, and shoot length was recorded from one experiment.

### Phylogenetic analysis with *Dyella* sp. A4

To determine phylogenetic relationships with other *Dyella* strains, the 16S rRNA gene sequence of *Dyella* sp. A4 was compared to sequences listed in GenBank using BLASTN. Sequences with the highest similarity along with more distantly related species within the *Dyella* genus and other genera were aligned with MUSCLE 5.1 (Edgar, 2004). A phylogenetic tree was constructed by the maximum likelihood method, based on Kimura’s two-parameter model with bootstrap analysis (1,000 replications) using PhyML 3.3.2 (Guindon et al., 2010).

### Inoculation of Arabidopsis with fluorescently-labeled *Dyella* sp. A4 in agar and microscopy

The localization of *Dyella* sp. A4 along the root was investigated by fluorescence microscopy after chromosomal integration of mScarlet-I using a mini-Tn7 suicide plasmid (pMRE-Tn7-155) (Schlechter et al., 2018). An overnight culture of *Dyella* sp. A4 was diluted in fresh 0.5x TSB medium and grown to an OD600 of 0.4-0.6. Cells were collected by centrifugation (8,000 g, 5 min), washed twice, and resuspended in 200 µl of 300 mM sucrose. A hundred nanograms of plasmid DNA was added to 50 µl of recipient cells, followed by electroporation with settings of 2.5 kV/cm, 25 μF, and 200 Ω. Afterwards, the cells were mixed with 800 µl of 0.5x TSB medium and allowed to regenerate at 28°C for 2-3 h. They were then incubated onto 0.5x TSB agar containing 50 µg/ml kanamycin for 4 days. Single colonies were re-streaked onto 0.5x TSB agar and validated by PCR. For inoculation of Arabidopsis with A4::Tn7-mScarlet, the same procedure as for the wild-type (WT) strain was followed. Colonization of the roots was visualized using a Leica M205 FA stereo fluorescence microscope with an ET mCherry filter (Leica Microsystems). To confirm fluorescence, an overnight culture was pelleted, washed, and mounted in 1% agarose for imaging under a Leica DM 5000B fluorescence microscope, equipped with a S-Orange (N2.1) filter cube at 100x magnification.

### Inoculation of Arabidopsis with *Dyella* sp. A4 in soil

*Dyella* sp. A4 was inoculated into 25 ml of deionized water at OD600 concentrations of 0.0002, 0.0004, and 0.0008. Single 5.5 cm pots containing gamma-irradiated alder swamp soil mixed with sand (1:3 v/v) were saturated with the bacterial treatments overnight. Arabidopsis seeds were sown the next day and shoot fresh weight was measured after 6 weeks of growth. For a second experiment using potting soil, *Dyella* sp. A4 was prepared at a final OD600 of 0.0002. Pots containing Pindstrup potting soil mixed with sand (1:3 v/v) were saturated overnight with either the bacterial solution or deionized water as a control. Shoot fresh weight was measured 24 days after sowing.

### Statistical analysis

Analyses were performed in R by a two-sided t-test, two-tailed Mann-Whitney *U*-test, or ANOVA followed by Tukey post hoc test to determine significance. A p-value < 0.05 was taken as statistically significant.

## Results

The root-associated microbiota of Arabidopsis plants grown in soil collected from an alder swamp in North Zealand, Denmark (Figure 1A) was analyzed and compared with the microbiota from bulk soil. Unconstrained Principal Coordinate Analysis (PCoA) of Bray-Curtis distances revealed a clustering of bacterial communities between root (i.e. endosphere and rhizoplane) and bulk soil (Figure S2A). Permutational multivariate analysis of variance (PERMANOVA) with Bray-Curtis distances indicated that 66.64% of total variance could be explained by compartment (*P* < 0.01) (Table S1). Furthermore, bacterial alpha-diversity based on the Shannon index was significantly higher in the bulk soil than in the root (Figure S2B). Comparison of bacterial community composition at the phylum level revealed an enrichment of *Bacteroidota, Pseudomonadota* and *Chloroflexota* and depletion of *Acidobacteriota, Bacillota, Nitrospirota* and *Planctomycetota* in the roots (Figure S2C and Table S2). At the ASV level, 134 ASVs were enriched and 144 ASVs were depleted in root samples (relative abundance > 0.1% in at least one sample, edgeR, generalized linear model, *P* < 0.05, FDR < 0.2; Table S3). The results indicate a filtering of bacterial community recruited from the bulk soil to the root.

**Figure 1.**
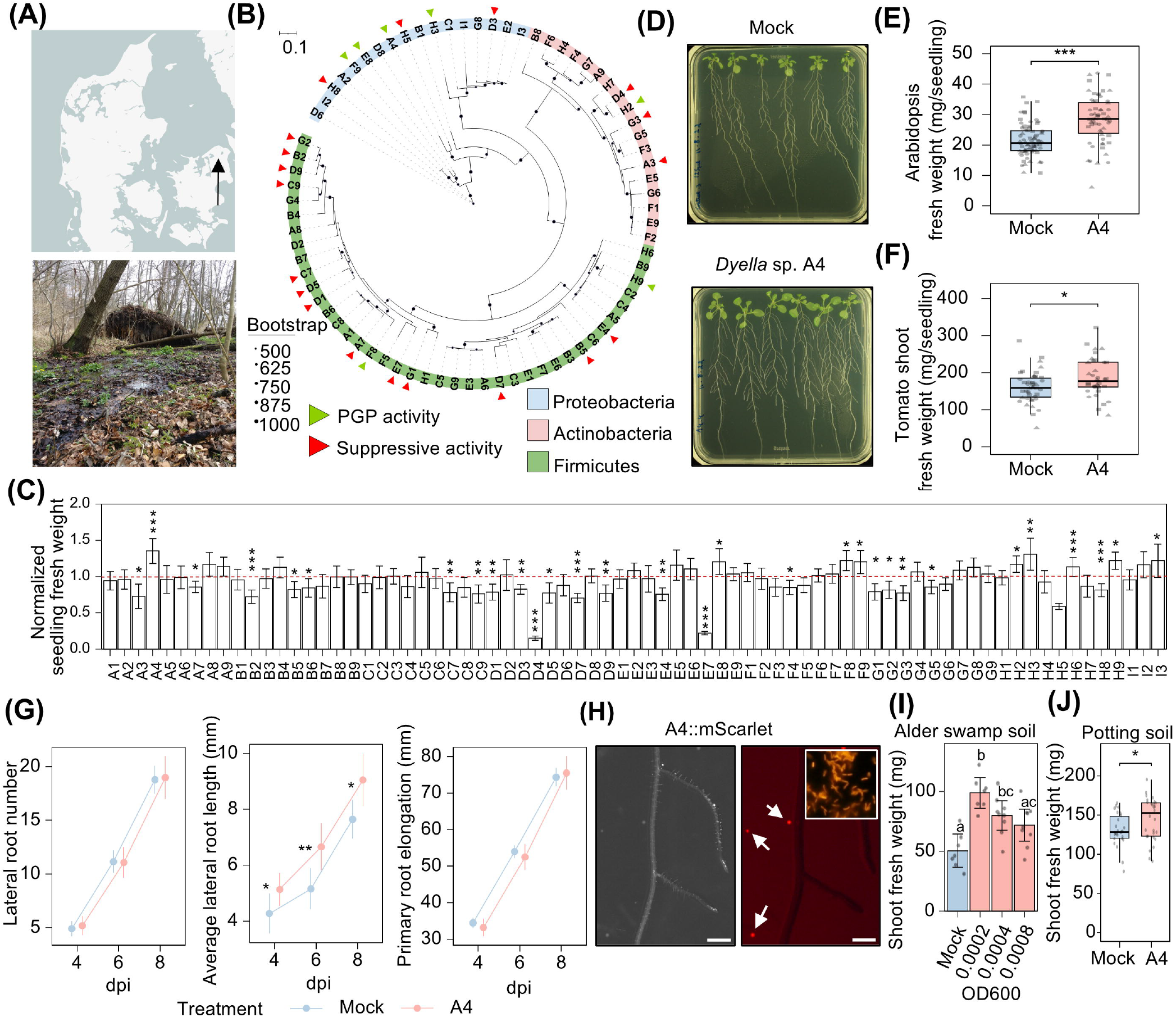
Identification of plant growth-promoting *Dyella* sp. A4 from Arabidopsis grown in an alder swamp soil collected in Danish nature. (A) Location (indicated by a black arrow) and biotope of the alder swamp soil sampling site in Denmark. (B) Consensus phylogenetic tree generated from the 16S rRNA gene sequences of 75 bacterial isolates. Tree tips are colored according to phylum and black circles represent bootstrap values based on 1,000 replicates. Green and red triangles depict strains with PGP and suppressive activity, respectively. (C) Fresh weight of Arabidopsis seedlings inoculated with the 75 strains in monoassociation normalized to axenic seedlings (indicated by a red dotted line). n = 15-24 seedlings per strain from one experiment. (D) Exemplary images of WT Arabidopsis growing in axenic agar (mock) or agar inoculated with *Dyella* sp. A4. (E, F) Biomass of Arabidopsis (E) and shoots of tomato (F) seedlings grown on agar inoculated with *Dyella* sp. A4 compared to axenic plates (mock). n = at least 59 plants per treatment in (E) and n = at least 37 plants per treatment in (F) pooled from three independent experiments (represented by different shapes). (G) Effect of *Dyella* sp. A4 on root development at four, six and eight days post inoculation (dpi). n = at least 36 plants per treatment x time point combination pooled from two independent experiments. (H) Stereo microscopy of Arabidopsis roots in agar at 10 dpi with A4::mScarlet under brightfield view (left) and fluorescent signal from mCherry filter (right). Arrows indicate fluorescent A4::mScarlet colonies in agar without association with the root. Scale bar represents 500 µm. Insert in the right panel shows fluorescence microscopy of A4::mScarlet cells from an overnight culture. (I) Effect of *Dyella* sp. A4 on shoot fresh weight of Arabidopsis grown in sterilized alder swamp soil when inoculated at different concentrations (OD600). n = 7-9 plants per treatment. (J) Effect of *Dyella* sp. A4 at OD600 of 0.0002 on shoot biomass of Arabidopsis grown in potting soil. n = 26-28 plants per treatment. Data represent mean ± 95% confidence interval in (C, G and I). Statistical significance was determined with two-tailed Mann-Whitney U-test in (C and G), by two-sided t-test in (E, F and J) and via ANOVA followed by Tukey post hoc test in I. *,** and *** indicate P < 0.05, 0.01, 0.001, respectively.

Towards the discovery of novel beneficial microbes for plant growth, we first isolated culturable bacteria from surface-sterilized roots of eight-week-old plants grown in the soil. A total of 290 purified isolates were obtained and identified by PCR amplification of ∼1500 bp from the 16S rRNA gene followed by Sanger sequencing. After removal of redundant isolates, we obtained a final culture collection comprised of 75 members (Figure 1B). The majority is represented by the *Firmicutes* phylum (53%), followed by *Actinobacteria* (25%) and *Proteobacteria* (22%). By sequence alignment of the V5-V7 region, we found that 72 strains of the 75 share > 97% 16S rRNA gene sequence identity to ASVs from the culture-independent analysis (Table S4), suggesting that the isolates are representative of the alder swamp soil-derived microbiota.

### Screening of isolates for plant growth promotion

To identify isolates with a PGP effect, we exposed six-day-old Arabidopsis seedlings to agar plates inoculated with each of the bacteria in monoassociation and measured total seedling biomass at 10 days post inoculation (dpi). Seven of the 75 strains significantly enhanced seedling fresh weight compared to the axenic controls (mock) (Figure 1B and 1C) and are found across all three phyla. Four belong to the *Proteobacteria* (*Massilia* sp. F9; *Paraburkholderia* sp. E8; *Dyella* sp. A4; *Paracoccus* sp. H3), two to the *Firmicutes* (*Paenibacillus* sp. F8; *Sporosarcina* sp. H9) and one to the *Actinobacteria* (*Arthrobacter* sp. H2). Among those, *Dyella* sp. A4 resulted in the highest increase in seedling biomass (27.6 ± 8.4) relative to the mock (19.9 ± 5.6 mg, Mann-Whitney *U*-test, *P* < 0.001). This growth promotion was reproducible across three independent experiments (Figure 1D and 1E) and *Dyella* sp. A4 was therefore selected for further analysis.

### Identification of a novel plant growth-promoting strain *Dyella* sp. A4

To search for candidate PGP-related genes in *Dyella* sp. A4, we sequenced its whole genome, performed de-novo assembly and predicted the coding sequences (CDSs). In total, 4373 CDSs were predicted and 4333 of those genes were complete genes. The predicted CDSs were annotated based on blast against the KEGG protein database. To annotate PGP-related genes, we curated a collection of PGP-related orthologs based on published literature, including genes involved in nitrogen fixation, phosphate solubilization and mineralization, siderophore biosynthesis and hormone modulation (auxin, cytokinin, giberellins, salicylic acid and ACC deaminase). The strain was identified as a new species when compared against currently available GTDB bacterial genome database and was found to harbor genes involved in organic phosphate mineralization, e.g. phosphodiesterase (K01126), phytase (K01093) and acid phosphatase (K09474), as well as phytohormone synthesis, e.g. metK (K00789) (ethylene), miaA (K00791) and miaB (K06168) (cytokinin), tyrB (K00832) and ALHD (K00128) (auxin), although none of the complete biosynthetic pathways of phytohormones were present (Table S5). We next tested whether the PGP potential could translate to another plant species. Inoculation of tomato seedlings with *Dyella* sp. A4 for 24 days in agar resulted in a higher shoot biomass (187.0 ± 51.5 mg) compared to the mock treatment (160.7 ± 42.4 mg, t-test, *P* < 0.05; Figure 1F), indicating that the stimulation of plant growth is not limited to the Brassicaceae family.

### *Dyella* sp. A4 can promote lateral root elongation

As the plasticity of the root system architecture is critical to maximize nutrient acquisition and plant growth, we monitored the root developmental response to *Dyella* sp. A4 at four, six and eight dpi. *Dyella* sp. A4**-**inoculated seedlings showed longer average lateral root lengths (5.1 ± 1.7 mm at four dpi, 6.7 ± 2.4 mm at six dpi, 9.0 ± 2.7 mm at eight dpi) compared to the mock (4.3 ± 2.4 mm at four dpi, 5.2 ± 2.5 mm at six dpi, 7.6 ± 2.3 mm at eight dpi), as determined by the Mann-Whitney *U*-test (*P* < 0.05; Figure 1G). By contrast, the number of lateral roots and primary root elongation were unaffected by the bacterial treatment. Auxin is a key regulator of root growth, we therefore monitored the auxin reporter DR5rev::GFP in the tip of primary roots after two and six days of inoculation. Consistent with the unchanged primary root elongation observed upon *Dyella* sp. A4 inoculation, the fluorescent pattern in the root tip showed no differences compared to the mock treatment (Figure S3).

To investigate the colonization pattern of *Dyella* sp. A4 in roots, we transferred six-day-old Arabidopsis seedlings onto agar inoculated with a fluorescently labeled version of the strain (A4::mScarlet), achieved through genome integration of mScarlet. Cells of A4::mScarlet expressed a strong fluorescent signal, however, we did not observe mScarlet signal on the rhizoplane or the root interior (Figure 1H). Instead, we noticed a few distinct fluorescent colonies in the agar medium. Comparison of the fresh weight of inoculated seedlings with those grown on mock-treated plates showed that A4::mScarlet retained the ability to significantly enhance plant biomass as previously demonstrated by the WT strain (Figure 1E and S4).

### *Dyella* sp. A4 promotes plant growth of soil-grown Arabidopsis

We investigated whether the PGP effect of *Dyella* sp. A4 translates to Arabidopsis grown in soil which is particularly important for agricultural application. Since the beneficial effect of microbial inoculants on plant growth is reported to be dose-dependent (Jensen et al., 2024; Suckstorff and Berg, 2003), we tested three inoculum concentrations of the bacterium. Six weeks after inoculation in the native, sterilized alder swamp soil, the lower and medium bacterial concentrations (OD600 of 0.0002 and 0.0004) stimulated an increase of shoot biomass (98.7 ± 14.0 mg and 80.0 ± 16.1 mg, respectively) compared to the mock treatment (50.4 ± 15.0 mg, ANOVA, *P* < 0.05; Figure 1I). The shoot fresh weight of plants inoculated with a higher bacterial concentration (OD600 of 0.0008) was not significantly different relative to the mock treatment, confirming a dose-dependent effect. The lower bacterial concentration (OD600 = 0.0002) also improved shoot fresh weight in potting soil (144.7 ± 27.6 mg) relative to control plants (130.9 ± 21.1 mg, t-test, *P* < 0.05; Figure 1J).

## Discussion

We isolated 75 different bacterial strains from surface-sterilized roots of Arabidopsis grown in an alder swamp soil and identified seven with PGP effects in agar (Figure 1C). *Dyella* sp. A4 showed the highest increase in seedling biomass and promoted the elongation of lateral roots without affecting the length of the primary root and lateral root numbers (Figure 1G). Another PGP rhizobacterium, *Bacillus subtilis strain* ALC_02, has been reported to stimulate lateral root length in Arabidopsis without affecting primary root growth or root number (Jensen et al., 2024). This strain produces the auxin indole-3-acetic acid (IAA), and its inoculation was shown to induce an accumulation of auxin in both the shoot and root tissues of Arabidopsis. Several *Dyella* strains previously isolated have been shown to produce IAA in vitro (Becerra-Castro et al., 2011; Palaniappan et al., 2010). While *Dyella* sp. A4 harbours the auxin biosynthesis-related genes tyrB and ALHD (Table S5), the complete pathway for IAA production was not identified in its genome. Consistently, multiple closely related *Dyella* strains (*D. japonica* XD53, *D. terrae* JS14-6, D. *kyungheensis* THB-B117) (Figure S5) tested negative for indole production, a precursor for auxin biosynthesis (Son et al., 2013; Weon et al., 2009; Xie and Yokota, 2005).

*Dyella* sp. A4 is able to promote plant growth and lateral root elongation without colonizing the root, possibly through emission of bioactive volatiles or the secretion of diffusible metabolites into the medium. Previously, cyclic peptides produced by *Pseudomonas* species, along with volatile organic compounds from *Bacillus* species, were reported to promote lateral root development and/or plant growth (Dutta et al., 2025; Ortiz-Castro et al., 2020; Zhang et al., 2007). A future task will be to characterize the molecular mechanisms employed by this strain to promote plant growth and modify root architecture.

In conclusion, we explored the microbial community isolated from plants grown in a soil collected from a Danish alder swamp for plant growth-promoting bacteria and identified seven PGP strains. Among them, a novel *Dyella* sp. A4 increased plant biomass in both agar and a complex soil environment. This is a case for the exploration of natural soils as a means to expand the toolbox of biostimulants for the green transition towards sustainable agriculture.

## Author contributions

D.X. and B.A.H. conceptualized and supervised the project. D.X., B.A.H. and J.R.P. Jespersen collected the alder swamp soil sample. L.D. isolated the bacterial culture collection and performed the inoculation experiments with support from J.L. and E.P. for inoculations. W.Z. conducted the 16S rRNA gene amplicon sequencing analysis and F.L. conducted the bacterial genome *de-novo* assembly, taxonomic classification and functional annotation. L.D. and D.X. wrote the paper based on an original draft prepared by L.D.. D.X., B.A.H., F.L., and Y.B. edited and reviewed the manuscript.

### Acknowledgements

We thank Ydun Kalsbeek-Hansen, Marlene Niedermayer, Silja Johanne Heilmann-Clausen and Hanna Rabe for laboratory assistance; the support staff at the Center for Advanced Bioimaging and growth facilities at Department of Plant and Environmental Sciences at University of Copenhagen for technical assistance; We acknowledge the financial support from Novo Nordisk foundation (NNF20OC0060824 to B.A.H), National Key Research and Development Projects (2022YFF1001800 to Y. B.) and Science Xplore Award (XPLORER-2023-1017 to Y.B.).

## Data availability

The 16S rRNA sequences of bacterial isolates are provided in the supplementary files S4 and S6. The raw genome sequencing data of *Dyella* sp. A4 are openly available in the zenodo database at DOI: <u>10.5281/zenodo.14673736</u>.

