## Supplementary figures for "Isolation of a novel plant growth-promoting *Dyella* sp. from a Danish natural soil"

[a]Present address: NIRAS A/S, Sortemosevej 19, 3450 Allerød, Denmark.

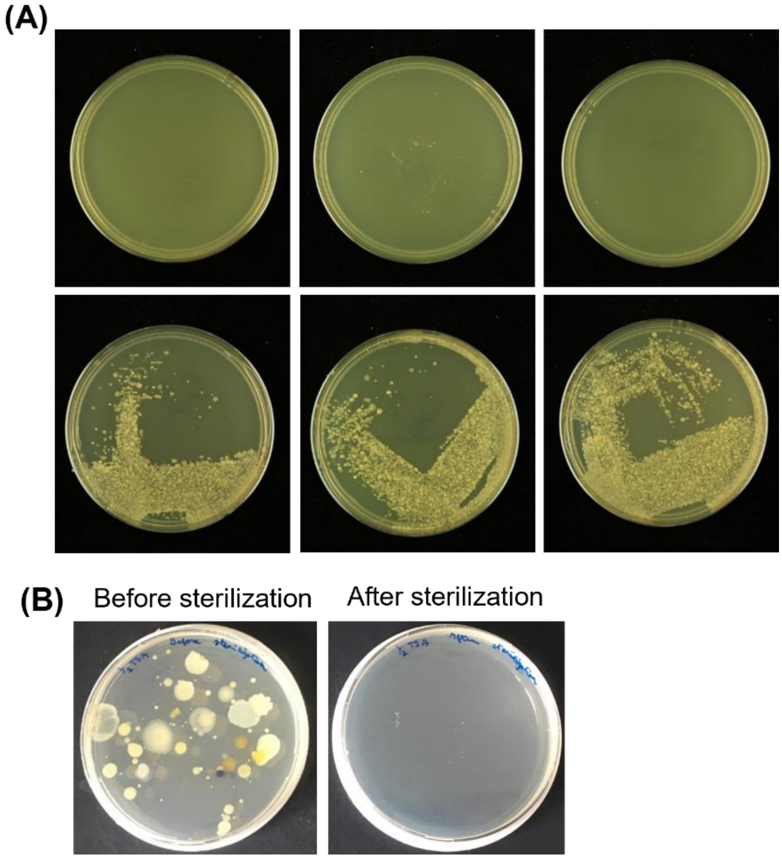


**Figure S1.** Verification of efficiency of sterilization. (A) Nutrient agar plates streaked with γ-irradiation-sterilized alder swamp soil (top) and non-sterilized alder swamp soil (bottom) in triplicates. (B) TSA plates streaked with water rinse from Arabidopsis roots before and after surface-sterilization.

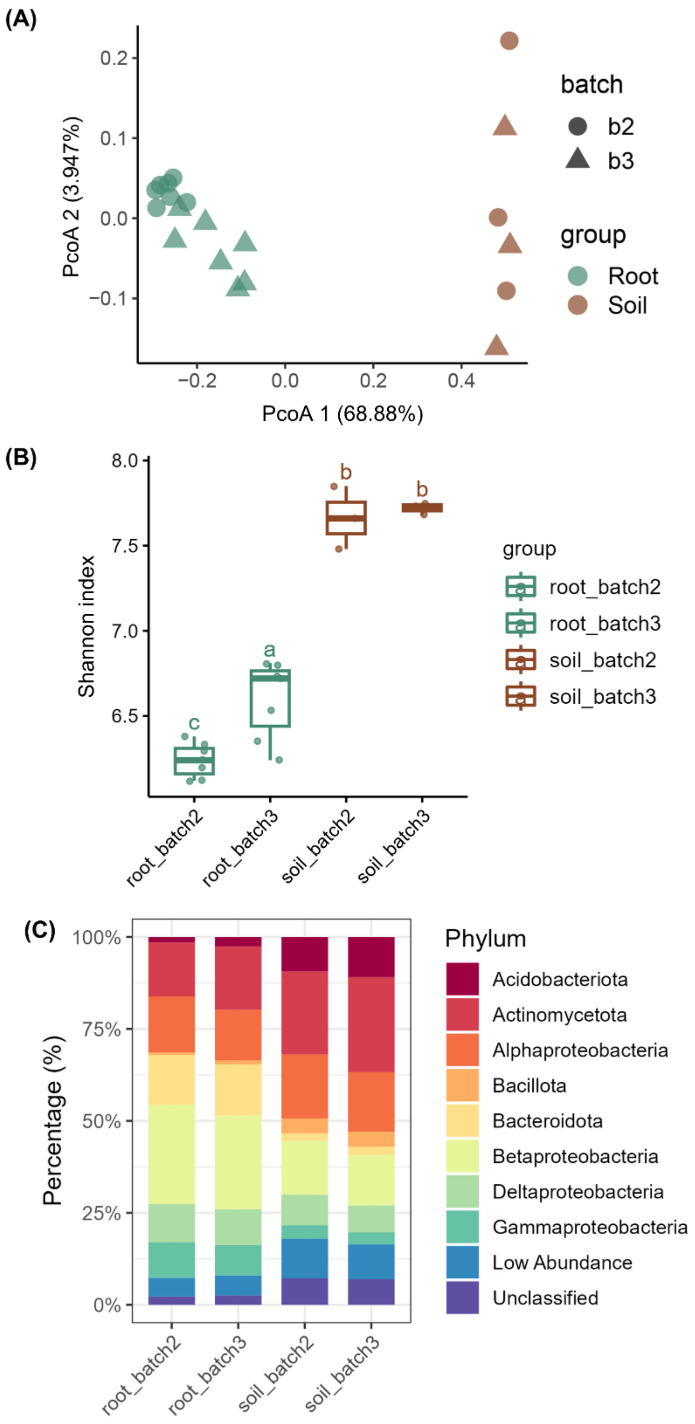


**Figure S2.** Differentiation of soil and root microbiota from alder swamp soil. (A, B) Principal Coordinate Analysis (PCoA) based on Bray-Curtis distances (A) and alpha-diversity based on Shannon index (B) for bacterial communities in root and soil samples. Different letters in (B) denote significantly different groups (Tukey HSD test, FDR adjusted $P < 0.05$). (C) Comparison of bacterial phyla relative abundance between soil and root samples. For each batch, n = 7 and 3 for root samples and soil samples, respectively.

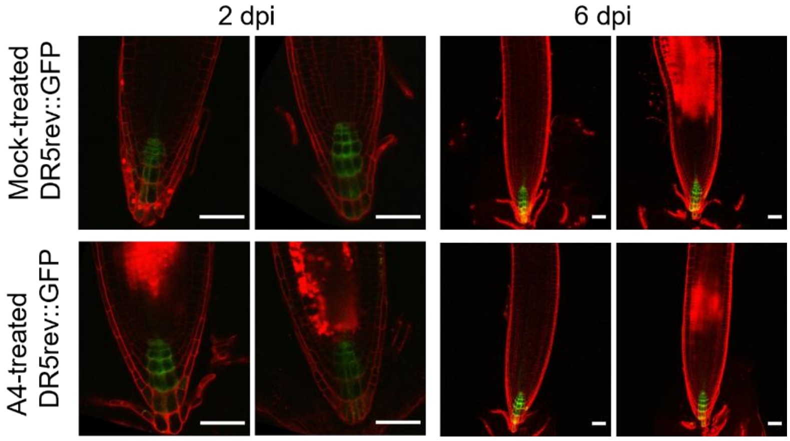


**Figure S3.** Monitoring the auxin response to *Dyella* sp. A4 in primary root tips. Confocal microscopy images of root tips of DR5rev::GFP auxin reporter grown on mock (top) or *Dyella* sp. A4-inoculated agar (bottom) at two and six dpi. Two different roots are shown per treatment and timepoint. Scale bars represent 50 µm.

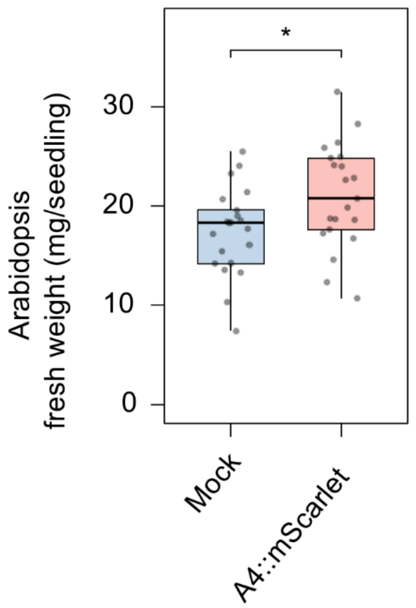


**Figure S4.** A4::mScarlet promotes Arabidopsis growth. Fresh weight of Arabidopsis seedlings grown on agar plates inoculated for ten days with A4::mScarlet compared to axenic plates (mock). n = 21 plants per treatment from one experiment. Significance was assessed via two-sided t-test (*indicates $P < 0.05$).

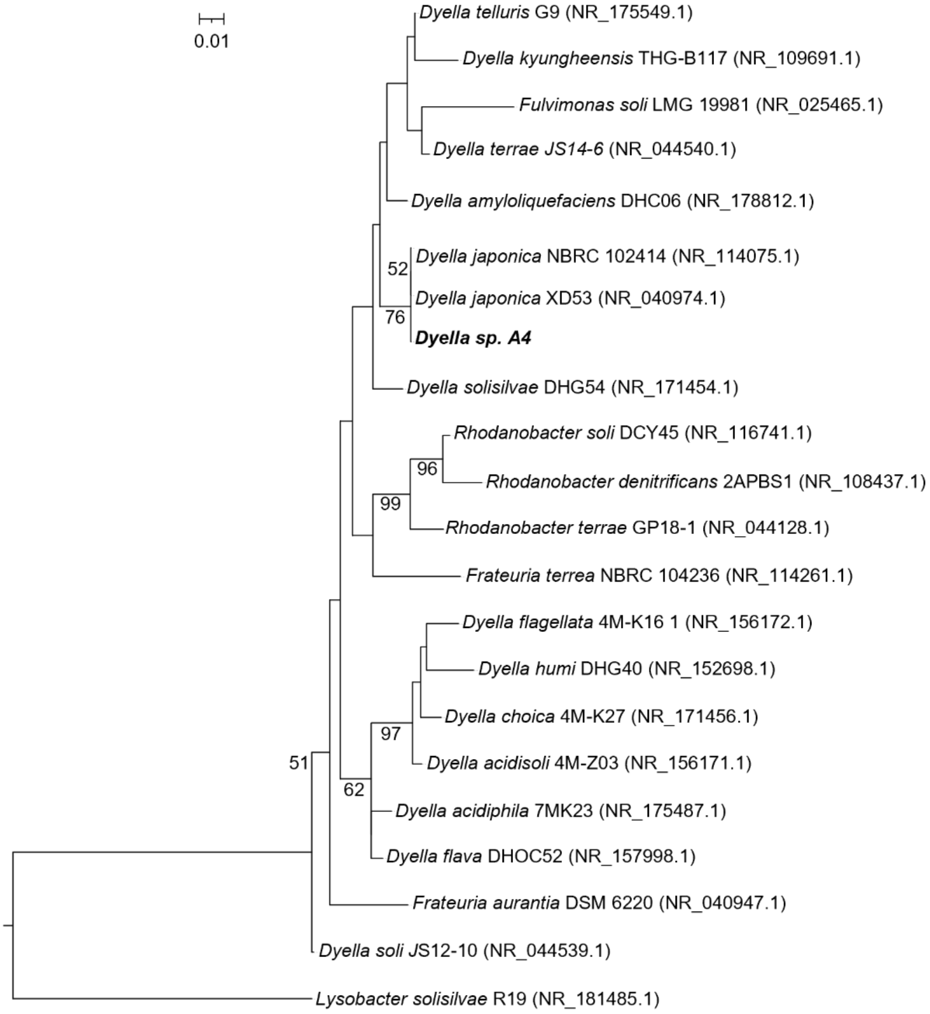


**Figure S5.** Maximum-likelihood phylogenetic tree based on 16S rRNA gene sequences showing the relationship of *Dyella* sp. A4 with other related species. *Lysobacter solisilvae* R19 was used as the outgroup. GenBank accession numbers of isolates are indicated after taxonomic names. Bootstrap percentages of 1,000 replicates are indicated for values greater than 50%.
